# Genomic Inbreeding and Selection Signatures analyses in the Doberman Pinscher breed

**DOI:** 10.64898/2026.06.04.730131

**Authors:** Henrique A. Mulim, Breno Fragomeni, Sophie Liu, Hinayah Rojas de Oliveira

## Abstract

The Doberman Pinscher population has undergone strong artificial selection for morphology and behavior, which can reduce genomic diversity and increase autozygosity. Here, we characterized the genome structure and identified selection signatures in Doberman Pinschers using complementary within- and between-population approaches. Genotypes from 3,226 Dobermans Dogs (Illumina CanineHD; 216,184 SNPs) provided by the Doberman Diversity Project were analyzed after purpose-specific quality control. Genomic inbreeding was quantified using four allele-frequency–based metrics and the runs of homozygosity (F_ROH_) approach. Selection signatures were detected using intrapopulation (i.e., Runs of Homozygosity—ROH; Integrated Haplotype Score—iHS; and Number of Segregating Sites by Length—nSL) and interpopulation methods (i.e., Fixation Index—FST; Cross-Population Extended Haplotype Homozygosity—XP-EHH; and Cross-Population Number of Segregating Sites by Length—XP-nSL) comparing the Doberman Pinscher breed to Labrador Retriever (n=237). Dobermans showed high overall inbreeding, with a mean F_ROH_ of 0.42 (range 0.22–0.68), whereas the allele-frequency–based inbreeding estimators had similar means (∼0.04). The partitioning of the ROH indicated high contributions from medium-to-long ROHs, consistent with recent inbreeding. The ROH scans identified 39,512 SNPs in ROH islands (≥50% frequency across individuals), with notable concentrations on CFA2, CFA3, and CFA31. Haplotype-based scans identified 2,820 candidate iHS SNPs and 2,173 candidate nSL SNPs (|score|>2). A common set of 310 SNPs was shared among ROH, iHS, and nSL, mapping near 279 genes that were mostly enriched for developmental pathways, particularly neurodevelopment and neuron-related cellular components. Between breeds, 349 highly differentiated SNPs were detected by FST, while XP-EHH and XP-nSL highlighted over 1,000 of Doberman-specific haplotype signals. A total of seven SNPs overlapped across FST, XP-EHH, and XP-nSL, which were located mainly on CFA8 (∼59.48–60.61 Mb) near the *KCNK10*, *SPATA7*, *PTPN21, NEGR1,* and *BTG1* genes. These genes are mainly linked to neural development and signaling, but *BTG1* has also been associated with cardiomyocyte cell-cycle regulation, and *KCNK10* with cardiac excitability and remodeling. Overall, the Doberman Pinscher breed exhibits high genome-wide autozygosity and levels of inbreeding. In addition, our results showed consistent, multi-method evidence of selection at loci associated with neurodevelopmental and regulatory pathways. These findings provide prioritized targets for follow-up studies that integrate phenotypes relevant to breed health and performance.

## Introduction

The Doberman Pinscher breed originated in the late 19th-century, in Germany, where Karl Friedrich Louis Dobermann, a tax collector, began developing the breed to serve as a loyal, protective companion and an effective guard dog during his rounds [1]. To achieve this, he crossed several breeds, including Rottweiler, Greyhound, and German Pinscher, producing a dog that combined agility with power [2]. Early in its history, the Doberman Pinscher became recognized for its intelligence, loyalty, and protective behavior, and it was widely adopted for working roles, including police and military services. The breed was formally registered with the American Kennel Club (AKC) in 1943, which further established its prominence as both a companion and working dog in the United States [1,2]

In terms of physical appearance, the Doberman Pinscher is a medium-to-large working breed with a lean, muscular, athletic build and a square, well-balanced outline (i.e., its body length is typically close to its height at the withers; [3]). The head is long, narrow, and wedge-shaped, and the coat is short, sleek, and close to the body. Common coat colors include black with rust markings, red (brown) with rust markings, blue with rust markings, and fawn with rust markings [4]. Overall, the breed’s posture and expression are generally described as alert, confident, and intelligent [5].

The strong and direct artificial selection over time has enabled the fixation of desirable morphological and behavioral traits across various dog breeds, making them more predictable in type and temperament [6]. These same breeding practices, however, can increase homozygosity not only for desirable alleles but also for harmful recessive variants, thereby contributing to the emergence and/or elevated frequency of inherited disorders within particular breed populations [7]. Accordingly, the careful management of genomic diversity is central to long-term genetic sustainability in dog breeds. Maintaining sufficient genetic diversity supports health, adaptability, and population viability, while also reducing the risks of inbreeding depression and breed-associated genetic diseases [8,9].

Characterizing the genomic consequences of such selective pressures requires analytical approaches capable of detecting selection footprints across the genome. Selection signature analysis is one of these approaches, designed to identify genomic regions influenced by natural or artificial selection through the detection of specific patterns such as extended regions with reduced genetic variation, long shared haplotypes, and marked differences in allele frequencies between populations or breeds [10,11]. In the context of a highly selected breed such as the Doberman Pinscher, such analyses can provide a deep understanding of genetic diversity and selection history, revealing regions of the genome that have been selected and exhibit reduced diversity.

The objectives of this study were to: (1) characterize the genomic structure of the Doberman Pinscher population; (2) detect selection signatures using three intrapopulation methods (i.e., Runs of Homozygosity—ROH; Integrated Haplotype Score—iHS; and Number of Segregating Sites by Length—nSL) and three interpopulation methods (i.e., Fixation Index—FST; Cross-Population Extended Haplotype Homozygosity—XP-EHH; and Cross-Population Number of Segregating Sites by Length—XP-nSL); and (3) perform gene annotation and functional analyses to identify candidate genes and metabolic pathways linked to genomic regions found in shared loci across the different selection-signature approaches.

## Material and Methods

### Genotypic Information

Genotypic information from 3,226 animals (1,489 males and 1,737 females), genotyped with the Illumina Canine HD genotyping array containing 216,184 SNP markers, was used in this study. The genotypic data were initially collected through routine genetic testing of collaborating dogs with the owners’ consent for the Doberman Diversity Project (https://www.dobermandiversityproject.org/), which were made available for this project.

Therefore, no animal care committee approval was necessary for this study, as all information required for this study was obtained from existing databases.

### Quality control

Different quality control procedures were used depending on the purpose of the analysis. For the analyses in which fixated markers were important (i.e., ROH, XP-EHH, and XP-nSL), a call-rate for animals and genotypes < 0.01 was used, and non-autosomal markers or markers without genomic position were removed. For the analyses that consider only segregating sites (i.e., FST, iHS, nSL), in addition to the initial quality control, markers with minor allele frequency (MAF) < 0.05 and deviating from Hardy-Weinberg equilibrium at the 10^-6^ level were also excluded from the dataset. In the end, 3,226 animals and 190,293 markers were used in the first set of analyses, and 3,226 animals and 61,910 markers were used for the second set of analyses.

### Intrapopulation analyses

#### Population structure and linkage disequilibrium

Population structure was evaluated using principal component analysis (PCA) implemented in PLINK v1.9 [12]. The PCA was computed from genome-wide SNP data using a variance-standardized genomic relationship structure, in which SNP covariances were scaled by their respective variances to avoid undue influence of markers with different allele-frequency variances.

Genome-wide linkage disequilibrium (LD) was estimated in PLINK software v1.9 [12] as the squared correlation (r2) between alleles at pairs of loci. To characterize LD decay with physical distance, marker pairs were grouped into distance bins, and mean r2 was calculated per bin: 10–100 kb using 10 kb intervals, and 100–1,000 kb using 100 kb intervals. Bins containing fewer than 50 SNP pairs were excluded to ensure stable mean LD estimates. The LD metric was computed using the PLINK options --r2 dprime-signed with-freqs --ld-window 100000 --ld-window-r2 0, which also outputs signed 𝐷′ and allele frequencies for comprehensive evaluation of LD patterns.

#### Inbreeding metrics

Five different inbreeding metrics were estimated using the genomic information. The first metric applied in this study was based on the additive genetic relationship (F_GRM_ - [13]), derived as:

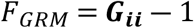

where

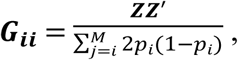

**G_ii_** is the genomic relationship matrix, **Z** is the nxM matrix of centered genotypes for n individuals and M markers, and p_i_ is the reference allele frequency in the population. The second method used in this study was based on the homozygous genotypes observed and expected (F_HOM1_ - [14]):

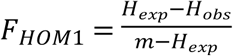

where H_exp_ is the expected value for homozygous genotypes, H_obs_ is the observed value for the homozygous genotypes, and m is the number of non-missing autosomal loci for each individual. The third method used was based on the excess of homozygosity, which was assessed by computing the mean per-SNP terms ratio instead of the ratios of sums (F_HOM2_ - Yang et al., 2011), i.e.:

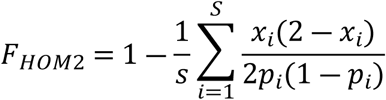

where xi ∈ {0,1,2} is the minor-allele count for the individual at SNP i, p_i_ is the minor allele frequency, and S is the number of SNPs.

As the three previous metrics strongly depend on allele frequency, we also tested another metric that reduces the impact of allele frequency on the estimate. This metric is based on the correlation between uniting gametes (F_UNI_ -[15] ), a direct estimation based on Wright’s definition of inbreeding [16]:

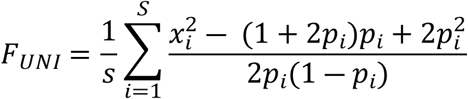

where x_i_ is the number of the reference allele copies of the i^th^ SNP, and p_i_ is the reference allele frequency in the population. The final metric used in this study was based on the sum of the individual lengths of ROH divided by the total length of the autosomal genome (F_ROH_ - [17]):

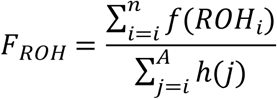

where (ROHi) is the total ROH length for individual i, n is the number of homozygous genomic regions in that individual, h(j) is the length of chromosome j, and A is the number of autosomal chromosomes (A=38). In addition, we calculated inbreeding within each ROH length class (<2 Mb, 2–4 Mb, 4–8 Mb, 8–16 Mb, and >16 Mb) as the sum of ROH lengths in that class divided by the total length of the autosomal dog genome. The first four genomic inbreeding coefficients were computed using PLINK v1.9 [12] while the last method was computed using an in-house R script. Correlations among the inbreeding metrics were evaluated using Pearson’s correlation coefficient [18], computed in R [19].

#### Runs of homozygosity

Runs of homozygosity were detected based on the windows approach using the PLINK v 1.9 software [12]. To reduce spurious ROH calls due to random homozygosity, we set the minimum number of markers required to define a ROH using the approach proposed by Lencz et al. [20] where the minimum number of markers (n_min_) is calculated as:

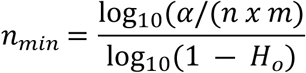

where α is the probability of observing a ROH by chance (set to 0.05), *n* is the number of individuals (3,226), *m* is the number of markers (190,293), and *Ho* is the population’s mean observed heterozygosity (0.32). Therefore, the minimum number of markers to define a ROH was 61 (--homozyg-snp 61). We further required ROH to be at least 500 kb long (--homozyg-kb 500), with a maximum inter-SNP density of 50 kb/SNP (--homozyg-density 50) and allowing gaps up to 200 kb (--homozyg-gap 200). To account for occasional genotyping errors, we allowed up to 1 heterozygous call within a ROH (--homozyg-het 1). The ROH were scanned using windows of 100 SNPs (--homozyg-window-snp 100), allowing up to 1 missing genotype per window (--homozyg-window-missing 1), and requiring at least 95% of overlapping windows supporting homozygosity (--homozyg-window-threshold 0.05). After identifying the ROHs, the segments were summarized in accordance with the length of the ROH and classified into five classes (<2, 2-4, 4-8, 8-16, and > 16Mb). Markers found in at least 50% of the cases inside of a ROH were considered candidate ROH islands for downstream annotation.

#### Integrated haplotype-based selection scans

Recent and ongoing positive selection was investigated using the integrated haplotype score (iHS) and number of segregating sites by length (nSL) statistics, which detect unusually long haplotypes associated with a given allele compared with the genome-wide expectation. Analyses were performed on phased haplotypes and run separately for each autosome using the software selscan v3.0.1 [21], then combined into genome-wide result files. The phased dataset used for iHS and nSL analyses included 3,226 animals and 61,910 SNPs, ensuring sufficient marker density and minimizing potential artifacts from low-quality or rare variants. Because a high-resolution genetic map was not available for all markers, we generated a proxy recombination map by assigning genetic positions under a constant recombination rate of 1 cM per Mb, using physical base-pair coordinates to derive genetic distance along each chromosome. The iHS and nSL were computed per chromosome using selscan [21], and the raw outputs were subsequently standardized (normalized) using the selscan norm utility with 20 allele-frequency bins to account for the dependence of these statistics on allele frequency. Candidate selection signals were defined as SNPs with absolute normalized iHS and nSL values greater than 2 [22–24], and these markers were retained for downstream visualization and annotation.

### Interpopulation analyses

#### Quality control in the Labrador population

For the interpopulation comparisons, we used data from 237 Labrador Retrievers genotyped with 166,171 SNP markers made public available by Campbell et al. [25]. After quality control and restricting to SNPs shared with the Doberman dataset, we generated two working datasets: one for fixation index (FST) analyses containing 47,688 common SNPs, and a second for cross-population (XP) analyses containing 144,855 common SNPs. The MAF and HWE filters were applied for the FST dataset because FST is estimated directly from allele-frequency differences, i.e., rare variants and markers with strong HWE deviations can produce unstable frequency estimates and inflate differentiation due to genotyping artifacts. In contrast, the XP analyses are haplotype-based and were performed on phased data; therefore, we did not additionally filter variants based on HWE for XP beyond the general genotype-level quality control and the requirement that variants are segregating.

#### Fixation Index

To reduce redundancy of SNP markers due to LD prior to estimating genetic differentiation, LD pruning was performed in PLINK v 1.9 [12] using a sliding-window approach (--indep-pairwise 50 5 0.8). This procedure evaluates LD within windows of 50 SNPs, shifts the window by 5 SNPs, and removes variants exceeding an LD threshold of r² = 0.8. Only SNPs retained after pruning were used in the dataset for the F_ST_ analysis (n=47,688). Genetic differentiation between the defined populations was quantified using Wright’s fixation index (F_ST_) as implemented in PLINK v 1.9 [12]:

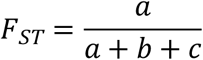

where *a* is the allele frequency variation among populations, *b* is the sampling variance, and *c* is the heterozygosity within populations. Markers with F_ST_ ≥ 0.80 were considered highly differentiated and were extracted for follow-up annotation.

#### Cross-population haplotype tests

To detect breed-specific recent selection, we performed cross-population haplotype scans using the selscan software [21], comparing Doberman (target population) against Labrador (reference population). Genotypes were phased per chromosome and population with the BEAGLE software [26], and phased VCFs files were used as direct inputs to selscan. The XP-EHH metric contrasts the integrated decay of extended haplotype homozygosity between populations and can be summarized as:

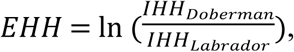

where IHH is the integrated haplotype homozygosity for each population, summarizing how long haplotypes remain homozygous (shared) around that marker. Thus, positive scores indicate unusually long, high-frequency haplotypes in Doberman relative to Labrador, while negative scores indicate stronger signals in Labrador.

Regarding the XP-nSL metric, the estimates will provide a complementary cross-population haplotype statistic, which is sensitive to sweeps across a range of allele frequencies. We quantified recent positive selection using the number of segregating sites by length (nSL) statistic and its cross-population extension (XP-nSL) as implemented in selscan [21]. The nSL is an EHH-based measure of haplotype length that replaces genetic distance with distance measured in numbers of SNPs, reducing sensitivity to variation in local recombination rates. The nSL score at the core site is then defined as the log-ratio of the two allele-specific integrals:

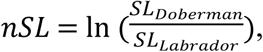

where *SL* is the allele-specific integrated haplotype homozygosity measured in SNP units. Raw XP-EHH and XP-nSL values were normalized with the selscan norm program using 20 bins to obtain standardized genome-wide scores for downstream thresholding and peak detection. Markers with absolute normalized XP-EHH and XP-nSL values ≥ 2 (i.e., |score| ≥ 2) were retained as candidate signals and were subsequently prioritized for downstream functional annotation.

### Functional Annotation

To biologically interpret candidate selection signals, we prioritized loci supported by multiple complementary approaches. Specifically, we defined two candidate marker sets: (i) SNPs shared among the intrapopulation analyses (ROH, iHS, and nSL), and (ii) SNPs shared between the interpopulation differentiation and cross-population haplotype tests (FST and XP statistics). These intersecting marker sets were carried forward for functional characterization, focusing on regions consistently identified across methods.

Functional annotation was performed in R using the GALLO package [27], version 2.0. Gene models were obtained from an Ensembl data file for the canine reference genome (CanFam3.1, [28]). Genes were retrieved using the gene-based method, considering a ±100 kb window around each marker. Gene-set enrichment was performed using the gprofiler2 package [29] available in R. Candidate gene symbols were tested for over-representation across functional categories using *Homo sapiens* as the reference organism to leverage curated functional resources and orthology-supported annotations. Enrichment significance was assessed using FDR multiple-testing correction, and terms with adjusted p<0.05 were retained as significant.

## Results

### Population Structure

Table 1 summarizes the genomic inbreeding coefficients for the Doberman Pinscher population. The average inbreeding for the first four metrics was the same (0.04), with values ranging from moderate negative to high positive inbreeding within the population. The metric with the highest inbreeding was the F_ROH_ metric (0.42), with the segments of 8-16Mb presenting the highest contribution to the value (0.12).

**Table 1.** Summary of different inbreeding coefficient metrics in the Doberman Pinscher population.

| Inbreeding | Min | Max | Mean | SD |
| --- | --- | --- | --- | --- |
| $F_{GRM}$ | -0.19 | 0.71 | 0.04 | 0.13 |
| $F_{HOM1}$ | -0.20 | 0.51 | 0.04 | 0.09 |
| $F_{HOM2}$ | -0.30 | 0.50 | 0.04 | 0.11 |
| $F_{UNI}$ | -0.10 | 0.49 | 0.04 | 0.07 |
| $F_{ROH}$ | 0.22 | 0.68 | 0.42 | 0.08 |
| $F_{<2Mb}$ | 0.02 | 0.11 | 0.05 | 0.01 |
| $F_{2-4Mb}$ | 0.02 | 0.15 | 0.06 | 0.01 |
| $F_{4-8Mb}$ | 0.04 | 0.20 | 0.10 | 0.02 |
| $F_{8-16Mb}$ | 0.01 | 0.23 | 0.12 | 0.03 |
| $F_{>16Mb}$ | 0.00 | 0.41 | 0.09 | 0.06 |

Min, minimum individual inbreeding coefficient; Max, maximum individual inbreeding coefficient; Mean, average of the inbreeding coefficient; SD, standard deviation of the inbreeding coefficients; $F_{GRM}$ , inbreeding coefficient based on the genotype additive variance; $F_{HOM1}$ , inbreeding coefficient based on the homozygous genotypes observed and expected; $F_{HOM2}$ , inbreeding coefficient based on homozygous genotypes; $F_{UNI}$ , inbreeding coefficient based on the correlation between uniting gametes; $F_{ROH}$ , inbreeding coefficient based on the length of the ROH's and the total length of the autosomal genome.

Fig 1 shows the correlation among the different metrics. In general, the correlation ranged from low to high and presented positive and negative correlations. Strong and positive correlations were found between F_HOM1_, F_HOM2_, F_UNI_, and F_ROH_. Also, the metrics F_HOM1_, F_HOM2_, and F_ROH_ had positive and high correlation with the long segments of ROH (F_8-16Mb_ and F_>16Mb_). Weak to moderate negative correlations were observed with the metrics of F_GRM_ and the other metrics, with F_UNI_ being the only metric to have a strong and positive correlation with F_GRM_ (0.63).

**Fig 1.**
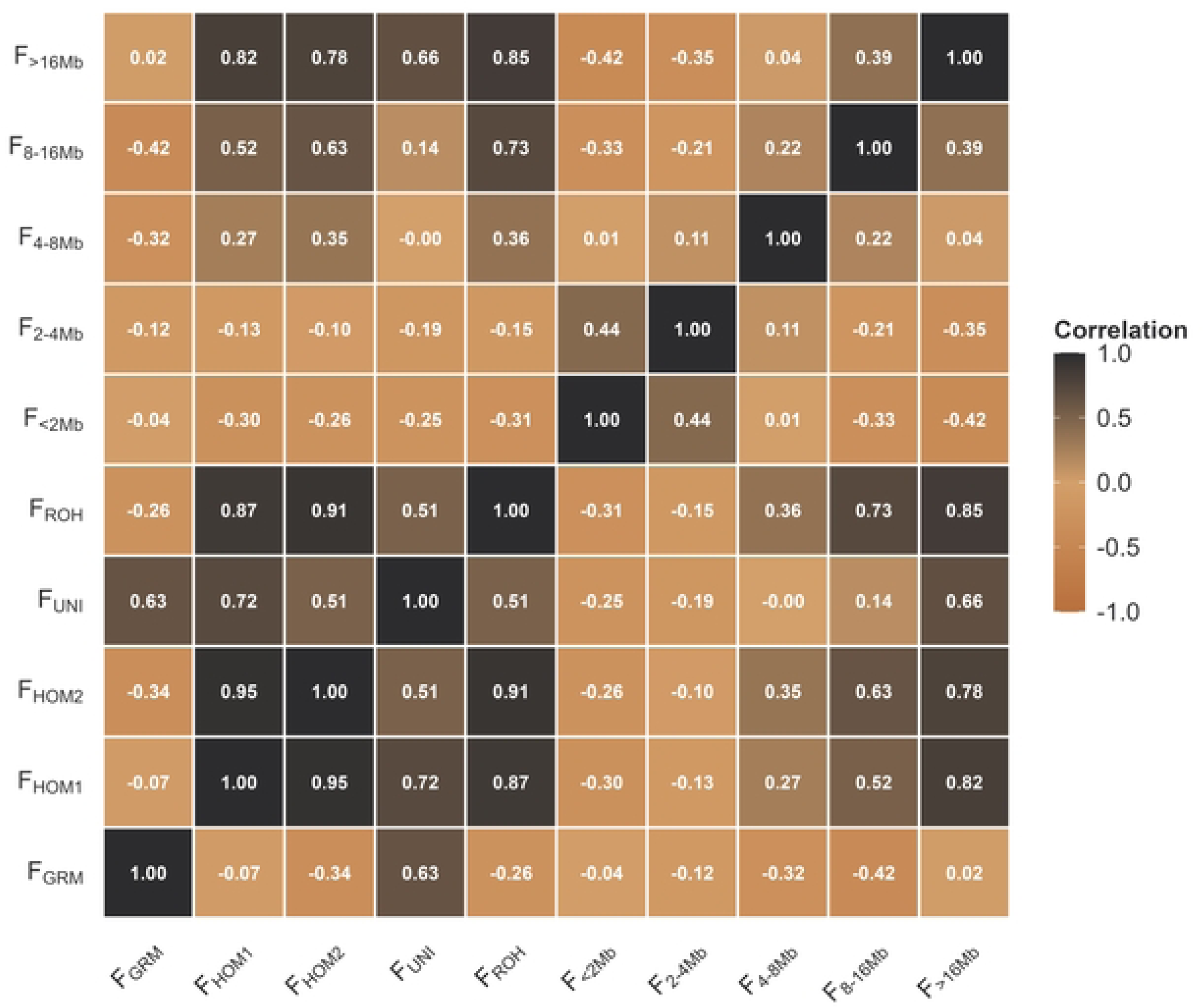
Correlation among the inbreeding metrics. F_GRM_: inbreeding coefficient based on the genotype additive variance; F_HOM1_: inbreeding coefficient based on the homozygous genotypes observed and expected; F_HOM2_: inbreeding coefficient based on homozygous genotypes; F_UNI_: inbreeding coefficient based on the correlation between uniting gametes; F_ROH_: inbreeding coefficient based on the length of the ROH’s and the total length of the autosomal genome

S1 Fig shows the LD decay and the plot of the first three PCAs for the Doberman Pinscher population. On average, the LD between the markers was higher at shorter distances (LD = 0.64 at 0.01kb) and decreased with increasing distance evaluated (LD = 0.20 at 1,000 kb). In the PCAs, the first three PCAs were responsible for explaining 7.35% of the variation among individuals. Fig 2 shows the distribution of the ROH per chromosome and the percentage of the chromosome covered by ROH.

**Fig 2.**
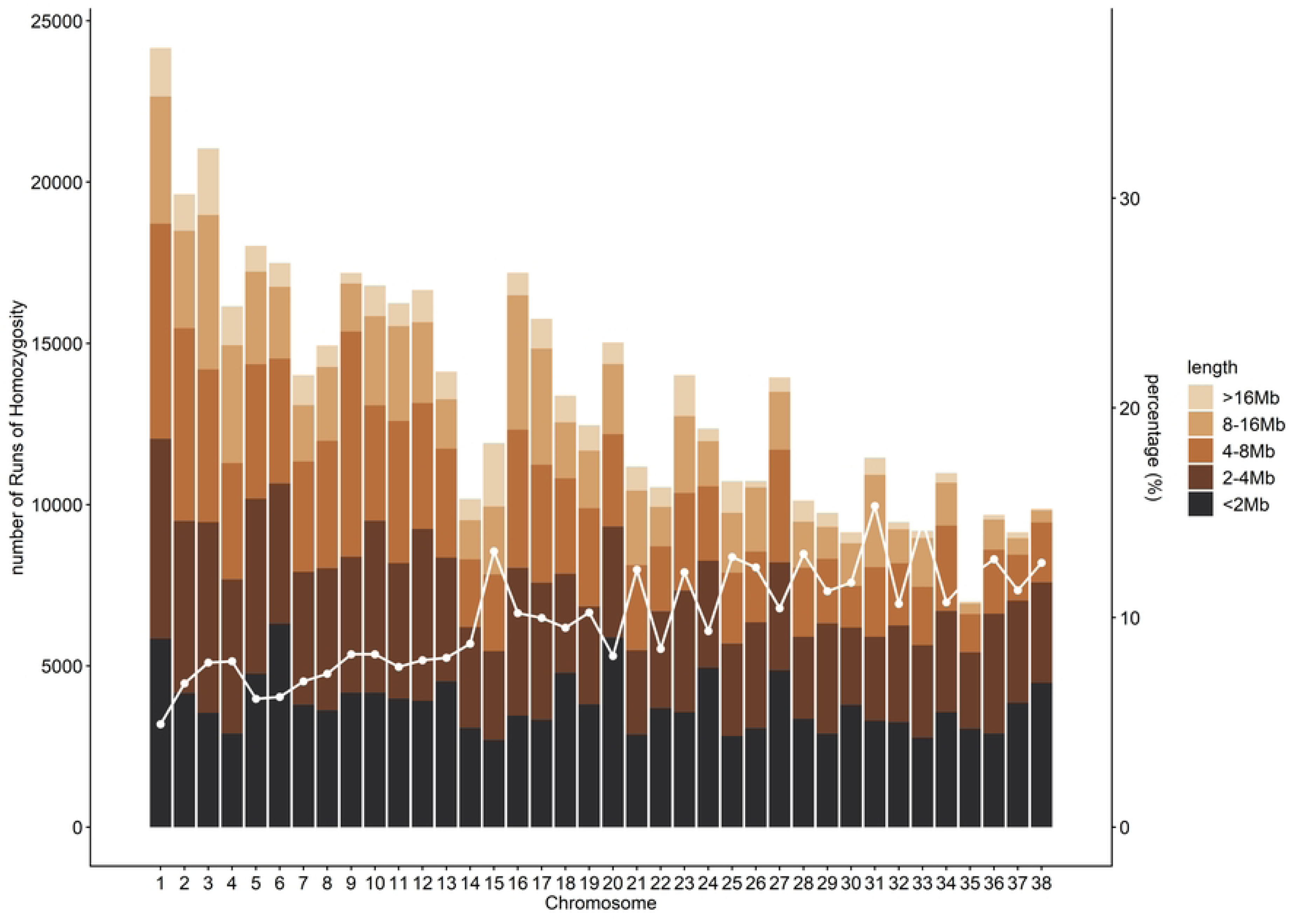
Distribution of Runs of Homozygosity per chromosome and length, and percentage of the chromosome covered by Runs of Homozygosity in the Doberman Pinscher population.

The chromosome with the highest number of ROHs was chromosome 1 (CFA1), and the chromosome with the highest coverage by the ROHs was CFA31 (∼15.65%). The majority of ROHs in the population correspond to segments of <2Mb (∼28.5%) and 2-4Mb (∼27.8%). Long segments of ROH (≥8Mb) corresponding to ∼20.6% of the total ROHs identified in the Doberman population, with ∼5.4% of the segments longer than 16Mb.

### Intrapopulation Selection Signature Methods

Fig 3 shows the Manhattan plots for the three intrapopulation selection-signature analyses performed in the Doberman Pinscher population, along with a Venn diagram showing the significant markers identified by each method and their overlap.

**Fig 3.**
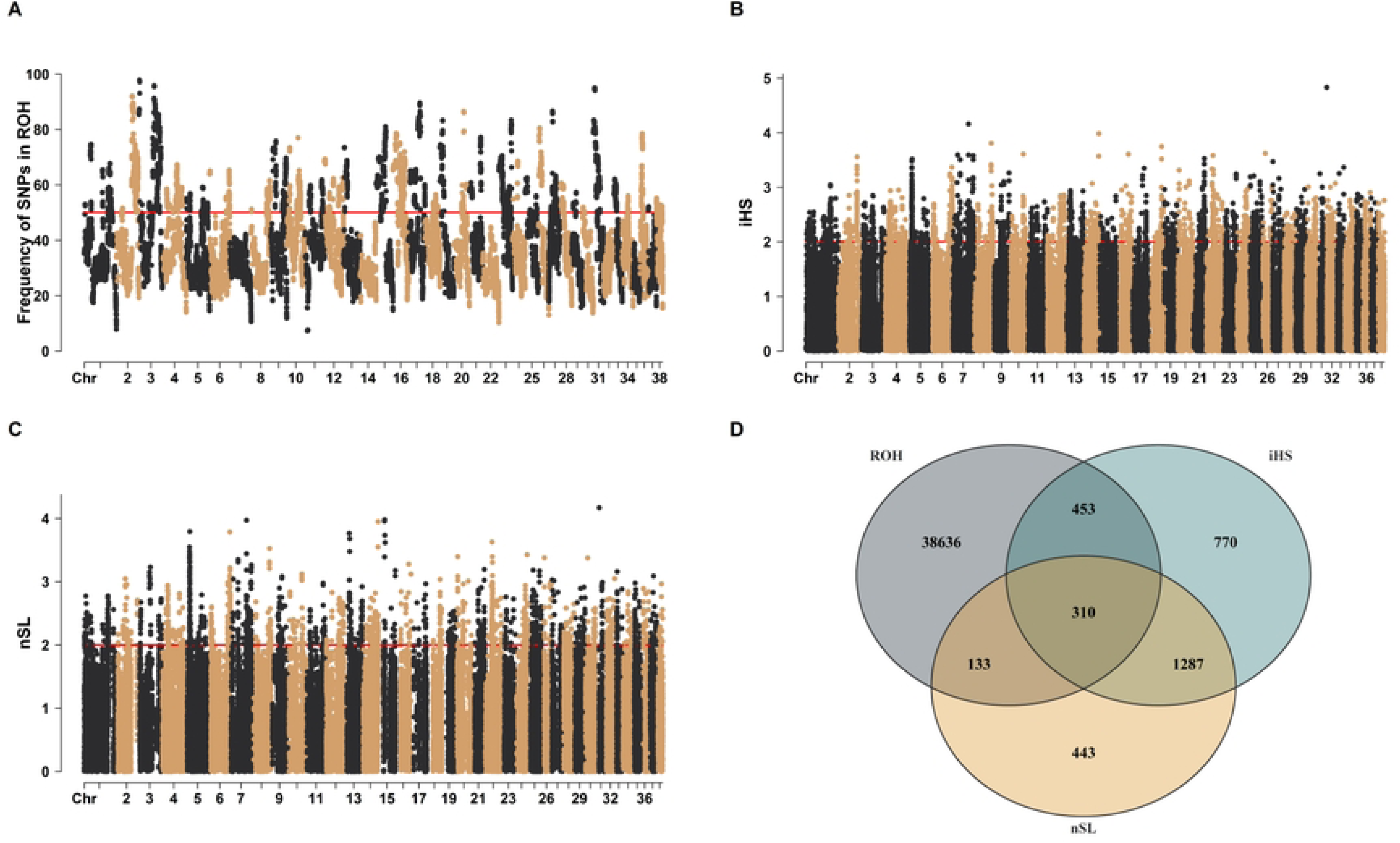
Genome-wide distribution and overlap of ROH islands and haplotype-based signatures of selection in Doberman Pinscher. (A) Frequency of SNPs within runs of homozygosity (ROH) across the genome. A 50% occurrence threshold was used to define ROH islands. (B) Manhattan plot of the integrated haplotype score (iHS) in the Doberman population. Points represent absolute normalized iHS values (|iHS|) for each SNP; loci with |iHS| > 2 were considered candidates for recent or ongoing positive selection. (C) Manhattan plot of the normalized number of segregating sites by length (nSL) values. Points represent absolute normalized nSL values (|nSL|) for each SNP; loci with |nSL| > 2 were considered candidates for recent positive selection based on unusually long haplotypes. (D) Venn diagram showing the overlap among candidate loci identified by ROH islands, iHS, and nSL analyses in the Doberman population.

In the ROH analysis, 39,512 markers were identified and located within ROH segments in more than 50% of the individuals. These markers were distributed across most chromosomes, except for CFA7, CFA14, CFA22, CFA32, and CFA35. For instance, 580 markers were mainly concentrated on CFA2, CFA3, and CFA31 (chromosomes that showed ROHs with highest concentration in the population). In the iHS analysis, 2,820 markers were identified with signs of recent ongoing selection (|iHS| > 2) in all autosomal chromosomes. From these, 763 markers were also identified in an ROH island. Regarding the sNL, the number of candidate markers for recent selection was 2,173. These markers were found across all autosomal chromosomes, and the number of markers in common with those in ROHs islands was 443, and with iHS was 1,597.

In total, 310 markers were shared across all three methods, providing strong evidence of selection signatures in these regions. These markers were located near genomic regions containing 279 annotated genes, including protein-coding genes (176), long non-coding RNAs (74), microRNAs (3), pseudogenes (4), small nucleolar RNAs (4), and small nuclear RNAs (18). Table 2 summarizes a subset of the genes located near markers identified by all three methods. The complete gene list is provided in S1 Table.

**Table 2.** Candidate genes located near genomic markers identified by all three methods of intrapopulation signatures of selection.

| CHR | START | END | ENSEMBL GENE ID | GENE NAME |
| --- | --- | --- | --- | --- |
| 1 | 92,898,274 | 92,978,260 | ENSCAFG00000002067 | *SLC1A1* |
| 1 | 93,142,635 | 93,435,997 | ENSCAFG00000002102 | *JAK2* |
| 5 | 14,503,902 | 14,509,113 | ENSCAFG00000029265 | *THY1* |
| 5 | 14,525,797 | 14,549,291 | ENSCAFG00000012042 | *USP2* |
| 5 | 14,564,935 | 14,705,257 | ENSCAFG00000012187 | *NLRX1* |
| 5 | 14,567,238 | 14,598,149 | ENSCAFG00000012079 | *MCAM* |
| 5 | 14,607,013 | 14,676,956 | ENSCAFG00000012100 | *CBL* |
| 5 | 14,795,750 | 14,807,262 | ENSCAFG00000012126 | *HYOU1* |
| 6 | 73,823,826 | 74,634,764 | ENSCAFG00000020411 | *NEGR1* |
| 8 | 59,484,328 | 59,612,750 | ENSCAFG00000017343 | *KCNK10* |
| 8 | 67,622,708 | 67,714,404 | ENSCAFG00000017809 | *BCL11B* |
| 12 | 1,341,191 | 1,354,584 | ENSCAFG00000000669 | *EHMT2* |
| 12 | 1,453,260 | 1,509,749 | ENSCAFG00000000751 | *TNXB* |
| 12 | 1,515,503 | 1,525,110 | ENSCAFG00000000761 | *ATF6B* |
| 18 | 47,635,788 | 47,912,117 | ENSCAFG00000010458 | *SHANK2* |
| 31 | 22,248,092 | 22,289,802 | ENSCAFG00000024758 | *ADAMTS5* |
| 33 | 7,266,685 | 7,418,106 | ENSCAFG00000009506 | *ABI3BP* |

CHR: canine chromosome (CFA); START and END: genomic coordinates (bp) based on the CanFam3.1 reference genome; ENSEMBL GENE ID: Ensembl gene identifier; GENE NAME: official gene symbol.

Table 3 shows the gene ontology terms enriched for biological process, cellular components, and molecular functions associated with the candidate genes located near genomic markers identified by all three methods of intrapopulation signatures of selection.

**Table 3.**
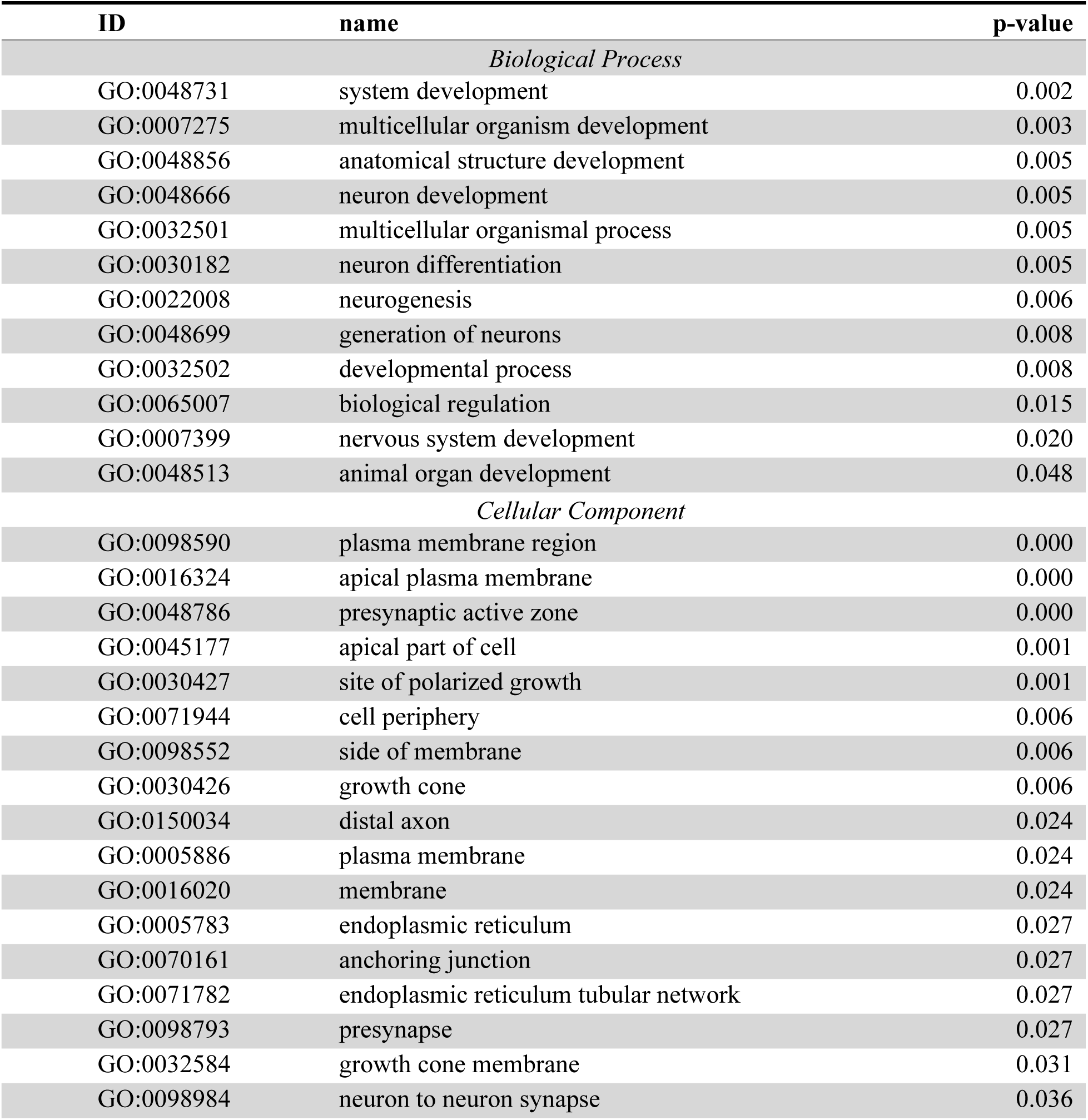
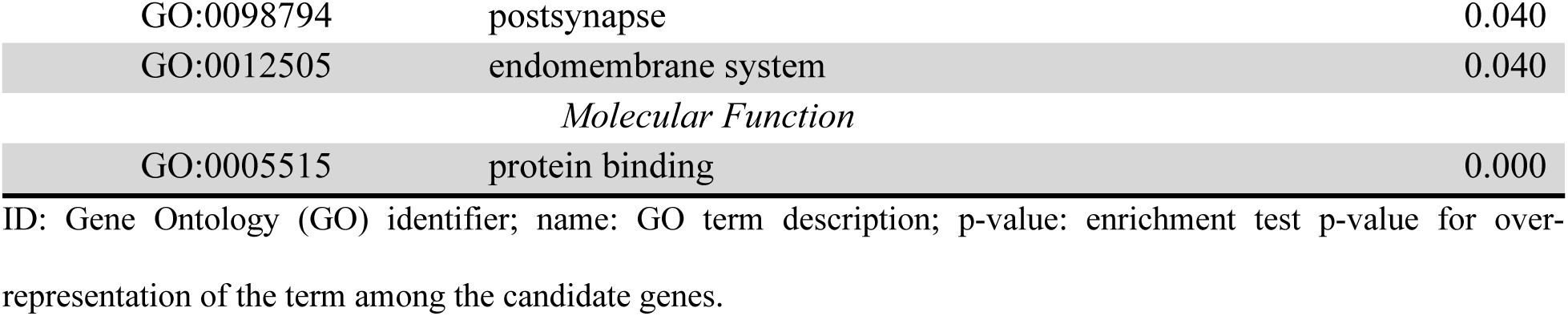
Gene Ontology terms enriched among genomic regions shared by all intrapopulation selection-signature methods.

| ID | name | p-value |
| --- | --- | --- |
| *Biological Process* |  |  |
| GO:0048731 | system development | 0.002 |
| GO:0007275 | multicellular organism development | 0.003 |
| GO:0048856 | anatomical structure development | 0.005 |
| GO:0048666 | neuron development | 0.005 |
| GO:0032501 | multicellular organismal process | 0.005 |
| GO:0030182 | neuron differentiation | 0.005 |
| GO:0022008 | neurogenesis | 0.006 |
| GO:0048699 | generation of neurons | 0.008 |
| GO:0032502 | developmental process | 0.008 |
| GO:0065007 | biological regulation | 0.015 |
| GO:0007399 | nervous system development | 0.020 |
| GO:0048513 | animal organ development | 0.048 |
| *Cellular Component* |  |  |
| GO:0098590 | plasma membrane region | 0.000 |
| GO:0016324 | apical plasma membrane | 0.000 |
| GO:0048786 | presynaptic active zone | 0.000 |
| GO:0045177 | apical part of cell | 0.001 |
| GO:0030427 | site of polarized growth | 0.001 |
| GO:0071944 | cell periphery | 0.006 |
| GO:0098552 | side of membrane | 0.006 |
| GO:0030426 | growth cone | 0.006 |
| GO:0150034 | distal axon | 0.024 |
| GO:0005886 | plasma membrane | 0.024 |
| GO:0016020 | membrane | 0.024 |
| GO:0005783 | endoplasmic reticulum | 0.027 |
| GO:0070161 | anchoring junction | 0.027 |
| GO:0071782 | endoplasmic reticulum tubular network | 0.027 |
| GO:0098793 | presynapse | 0.027 |
| GO:0032584 | growth cone membrane | 0.031 |
| GO:0098984 | neuron to neuron synapse | 0.036 |
| GO:0098794 | postsynapse | 0.040 |
| GO:0012505 | endomembrane system | 0.040 |
| *Molecular Function* |  |  |
| GO:0005515 | protein binding | 0.000 |

ID: Gene Ontology (GO) identifier; name: GO term description; p-value: enrichment test p-value for over-representation of the term among the candidate genes.

Functional enrichment of genes located near markers shared across all intrapopulation selection-signature methods revealed an over-representation of developmental pathways, particularly related to neurodevelopment. The most significant biological-process terms were related to system and organismal development, including neuron development, differentiation, and neurogenesis (p ≤ 0.020). Consistently, enriched cellular-component terms highlighted plasma-membrane and neuron-specific structures, including presynaptic active zone, growth cone, distal axon, and synaptic compartments (p ≤ 0.040). At the molecular-function level, protein binding was the predominant enriched term (p < 0.001).

### Interpopulation Selection Signature Methods

Fig 4 shows the Manhattan plot for the FST and cross-population analysis comparing the Doberman population to the Labrador Retriever population, along with the Veen diagram comparing the significant markers identified in each one of the analyses.

**Figure 4.**
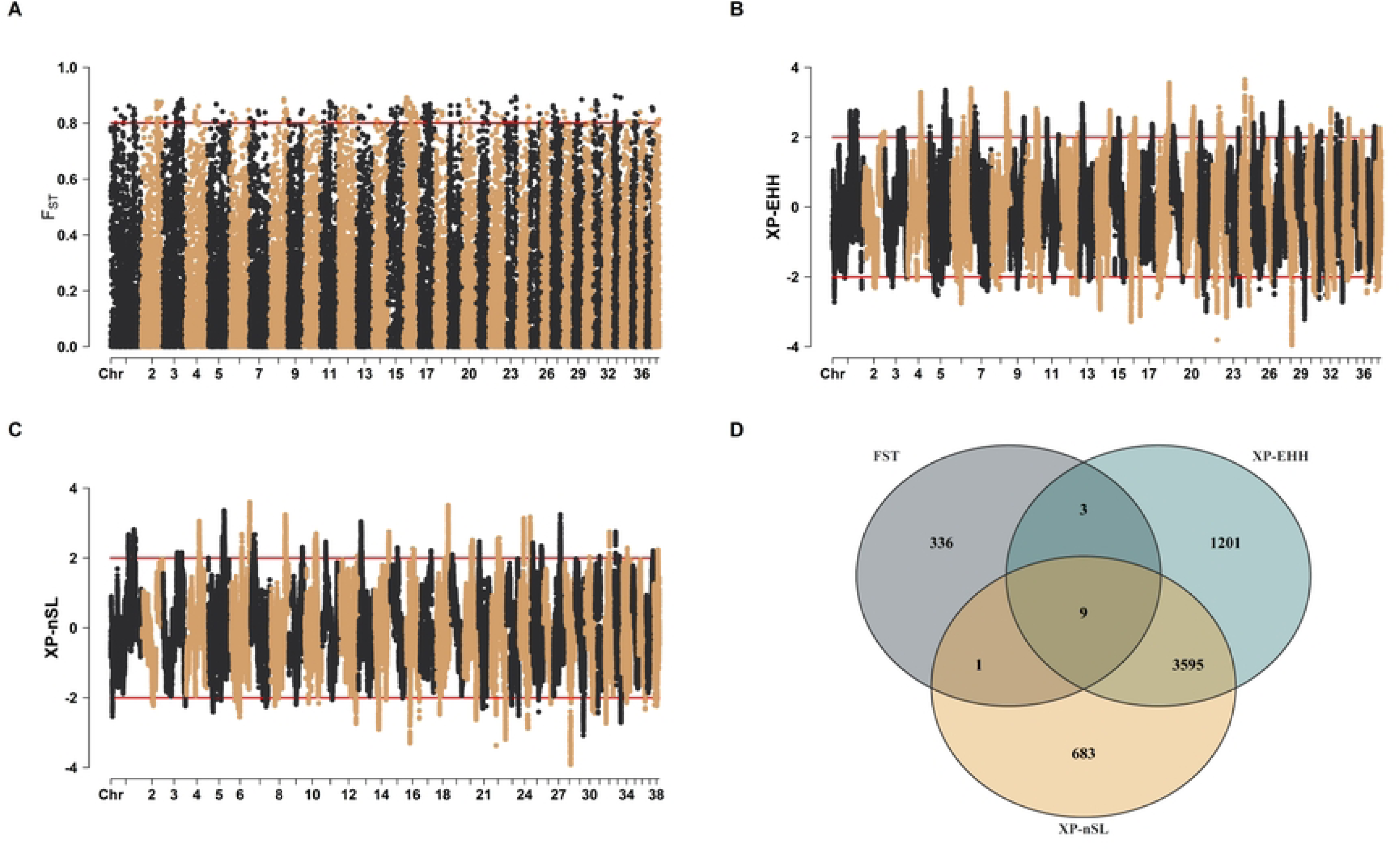
Genome-wide distribution and overlap of interpopulation selection signatures (FST, XP-EHH, and XP-nSL) between the Doberman Pinscher and the Labrador Retriever. (A) Genome-wide distribution of FST between Doberman and Labrador populations. SNPs within the upper 20% of the empirical FST distribution (≥80th percentile) were considered candidates for population-differentiated selection. (B) Cross-population extended haplotype homozygosity (XP-EHH) comparing Doberman (target) and Labrador (reference) populations. Horizontal red lines at XP-EHH = +2 and XP-EHH = −2 denote thresholds for strong population-specific selection. (C) Signed Cross-Population Number of Segregating Sites by Length (XP-nSL) analysis contrasting Doberman and Labrador populations. Horizontal red lines at XP-nSL = +2 and XP-nSL = −2 indicate thresholds for strong population-specific selection. (D) Venn diagram showing the overlap among inter-population selection signals identified using FST, XP-EHH, and XP-nSL analyses comparing Doberman (target) and Labrador (reference) populations.

For the FST analysis, 349 markers showed a high differentiation among the populations. These markers are present in all autosomal chromosomes, with CFA16 (32), CFA3 (28), CFA2 (25), and CFA11 (35) being the chromosomes that present the highest number of markers exceeding the threshold. For the XP-EHH, 3,101 markers had a differentiation from the reference population (high positive values). These markers were present in almost all autosomal chromosomes, except in CFA26 and CFA28. Ten markers were found in common among the XP-EHH and the FST metric. For the XP-nSL method, 2,832 markers showed a high differentiation from the reference population. These markers are distributed along almost all autosomal chromosomes except CFA2, CFA23, CFA26, CFA28, and CFA36. The number of markers in common between the XP-nSL method and FST was 8, and between XP-nSL and XP-EHH was 2,441.

Seven markers were found to be common across all three methods. These markers are located close to regions that encode 13 genes, as shown in Table 4. From these, 9 were protein-coding and 4 long non-coding RNAs (lncRNAs).

**Table 4.** Candidate genes near genomic markers identified by all three methods of interpopulation signatures of selection.

| CHR | START POS | END POS | GENE ID | GENE NAME | GENE BIOTYPE |
| --- | --- | --- | --- | --- | --- |
| 1 | 93,846,269 | 93,957,359 | ENSCAFG00000002131 | *ERMP1* | Protein coding |
| 1 | 94,096,954 | 94,103,734 | ENSCAFG00000002137 |  | Protein coding |
| 6 | 73,823,826 | 74,634,764 | ENSCAFG00000020411 | *NEGR1* | Protein coding |
| 8 | 59,484,328 | 59,612,750 | ENSCAFG00000017343 | *KCNK10* | Protein coding |
| 8 | 59,565,623 | 59,565,877 | ENSCAFG00000041412 |  | Protein coding |
| 8 | 59,658,365 | 59,697,318 | ENSCAFG00000017354 | *SPATA7* | Protein coding |
| 8 | 59,708,972 | 59,782,459 | ENSCAFG00000017377 | *PTPN21* | Protein coding |
| 8 | 60,260,126 | 60,275,034 | ENSCAFG00000043039 |  | lncRNA |
| 8 | 60,275,952 | 60,277,947 | ENSCAFG00000047705 |  | lncRNA |
| 8 | 60,288,179 | 60,289,810 | ENSCAFG00000048045 |  | lncRNA |
| 8 | 60,331,172 | 60,605,478 | ENSCAFG00000017486 |  | Protein coding |
| 15 | 32,647,472 | 32,647,762 | ENSCAFG00000006153 | *BTG1* | Protein coding |
| 15 | 32,657,752 | 32,666,640 | ENSCAFG00000046390 |  | lncRNA |

CHR: canine chromosome (CFA); START and END: genomic coordinates (bp) based on the CanFam3.1 reference
genome; ENSEMBL GENE ID: Ensembl gene identifier; GENE NAME: official gene symbol; GENE BIOTYPE:
gene classification based on the type of RNA transcript produced.

Most markers mapped to a single region on CFA8 (∼59.48–60.61 Mb), which included multiple protein-coding genes (*KCNK10*, *SPATA7*, *PTPN21*) and several nearby lncRNAs. Additional markers were detected on CFA1 (*ERMP1* and one unannotated protein-coding locus), CFA6 (*NEGR1*), and CFA15 (*BTG1* and one lncRNA).

## Discussion

### Population Structure

The main goal of this study was to characterize population structure and identify signatures of selection in Doberman Pinschers using multiple complementary approaches. Across the first four genomic inbreeding coefficients used in this study, mean inbreeding was similar (0.04; Table 1). However, as the methods capture different aspects of autozygosity and are scaled relative to allele-frequency expectations, distinct ranges (including the presence of both negative and positive values) were identified for the different metrics. For instance, the F_GRM_ quantifies an individual’s homozygosity relative to the sample mean using allele frequencies to define expected genotype variance, such that positive values indicate individuals that are more homozygous (i.e., more related) than the population average, and negative values indicate less homozygosity than expected [13]. The homozygosity-based estimators (F_HOM1_ and F_HOM2_) summarize deviations in observed homozygosity from the expectation under Hardy–Weinberg proportions given the population allele frequencies, thereby reflecting excess homozygosity consistent with identity-by-descent. In this context, positive values indicate excess homozygosity, whereas negative values indicate an excess of heterozygosity relative to expectation [14,15]. The F_UNI_ aims to measure autozygosity with reduced dependence on allele frequency, with the scaling focusing on the correlation between the two gametes that formed the individual; i.e., positive values indicate more correlated gametes (greater autozygosity), whereas negative values indicate less correlation [15,30].

Overall, frequency-based estimators (F_GRM_, F_HOM1_, F_HOM2_, F_UNI_) showed comparable means and included both negative and positive values (Table 1), consistent with their dependence on allele-frequency centering within the study sample. In contrast, ROH-based inbreeding was substantially higher (mean F_ROH_ = 0.42; range 0.22–0.68), indicating extensive autozygosity in this population. An advantage of the ROH method is that it measures the proportion of the genome that is homozygous and, by grouping ROH by length, gives an estimate of when inbreeding occurred in the population [31]. This feature makes the method useful for population management, because not all inbreeding is harmful [32,33]. Knowing when inbreeding events occurred helps reconstruct the population’s history. Partitioning F_ROH_ by ROH length showed that most autozygosity was attributed to medium-to-long segments (4–16 Mb), consistent with recent inbreeding, whereas the >16 Mb class exhibited high variability (maximum 0.41), suggesting that a subset of individuals experienced very recent consanguineous matings.

Across metrics, the strong concordance between F_HOM_-based measures and F_ROH_ indicates that the main inbreeding signal is robust and not driven by a single estimator (Fig 1). An interesting point to observe is that most metrics have a high correlation with the more recent types of inbreeding (F_8-16Mb_ and F_>16Mb_). This suggests that, in this population, the majority of the allele-frequency drive estimator captures more recent inbreeding than older events in the population. By contrast, weaker or negative associations between F_GRM_ and other estimators were observed. This might be due to the greater sensitivity of GRM-based inbreeding to allele-frequency centering and the chosen reference frequencies, which can shift the scale (and occasionally the sign) in structured or relatively homogeneous samples [34,35].

Recent inbreeding events in the population are also evident in the distribution of ROH across the genome (Fig 2). Long segments of homozygosity were detected in all autosomal chromosomes evaluated, corresponding to around 20.6% of the ROHs identified in the genome of the Doberman Pinscher population. Based on the length of the segments, it is estimated that consanguineous mating has happened in the population around 3 to 5 generations ago [31,36].

### Intrapopulation Selection Signatures Methods

We used three complementary methods to detect within-population signatures of selection in the Doberman Pinscher (Fig 3). First, ROH islands (Fig 3A) highlight genomic regions that are repeatedly homozygous across many individuals. Such patterns can emerge when strong and/or sustained selection increases the frequency of shared haplotypes, thereby reducing local genetic diversity [37]. By contrast, iHS (Fig 3B) is tailored to detect recent, incomplete selective sweeps. It does so by comparing haplotype homozygosity surrounding the ancestral versus derived allele at each SNP [22]. Finally, nSL (Fig 3C) is conceptually similar to iHS, but it measures haplotype length using the number of segregating sites rather than genetic distance. This reduces sensitivity to recombination-rate heterogeneity and can improve performance in regions where recombination varies across the genome [38]. Taken together, these approaches provide evidence across different time scales and genomic contexts: ROH islands are more consistent with longer-term or recurrent selection that increases autozygosity, while iHS and nSL are better at capturing sweeps that are still in progress and have not reached fixation.

Across all three methods, we found 310 SNPs that overlapped between ROH islands, iHS, and nSL, providing convergent support for loci under selection in this population (Fig 3D). This three-way overlap is notable because each method relies on different information (autozygosity versus haplotype structure) and different null expectations. Concordant signals are therefore less likely to reflect method-specific artifacts and more likely to represent true targets of selection or tightly linked variants. Genes near these shared markers point to biological themes involving extracellular matrix (ECM) remodeling, cellular stress responses, and vascular/cardiac biology, with additional connections to neurodevelopmental processes. These categories are plausible in Doberman Pinschers, given the breed’s history of intense artificial selection and the importance of cardiovascular and connective-tissue traits to breed health and performance [1].

Several overlapping candidates are consistent with an ECM and fibrosis-related signal. The *THY1* gene encodes a cell-surface glycoprotein involved in cell adhesion and stromal-cell biology, with established roles in wound healing and fibrotic remodeling, and it is widely used as a marker for particular fibroblast subpopulations across tissues [39–41]. In cardiac settings, *THY1*-positive fibroblast populations have been associated with fibrosis and reduced contractile function in experimental and translational work [42]. The *ADAMTS5* gene, which encodes a secreted metalloprotease (aggrecanase-2), cleaves ECM proteoglycans and is best known for its role in cartilage turnover and osteoarthritis-related degeneration [43,44]. While its strongest associations are musculoskeletal, *ADAMTS*-family proteases more broadly contribute to ECM remodeling, making *ADAMTS5* relevant if selection is acting on matrix dynamics across tissues [45]. The *ABI3BP* encodes an ECM-associated protein expressed in cardiovascular tissues and has been linked to cell–ECM interactions and remodeling phenotypes, supporting a potential role in structural maintenance and cardiovascular function [46]. The *TNXB* gene, a tenascin-family ECM gene, is also consistent with selection influencing connective-tissue architecture; although its direct link to specific Doberman Pinscher traits may be indirect, tenascin biology is often discussed in relation to tissue integrity and cardiovascular/connective-tissue pathology [47,48].

A second set of overlapping candidates fits stress-response and protein homeostasis pathways, which are repeatedly implicated in cardiac disease and hypertrophic remodeling. The *JAK2* gene is a key component of JAK/STAT signaling and is broadly involved in inflammatory and stress signaling, with extensive experimental support linking this pathway to cardiac hypertrophy and heart failure biology [49]. The genes *HYOU1* and *ATF6B* map to the unfolded protein response and endoplasmic reticulum stress pathways, which are activated under ischemic and hypertrophic stress and can influence cardiomyocyte survival and remodeling outcomes [50].

In line with this theme, *USP2* (a deubiquitinase) and *CBL* (an E3 ubiquitin ligase) connect to ubiquitin-mediated regulation of signaling and proteostasis; both have been reported in studies involving oxidative stress and hypertrophic or heart failure–related phenotypes, depending on the system and context [51,52]. The *NLRX1* gene, a mitochondrial innate immune receptor, further supports a mitochondria–stress axis, as it has been implicated in regulating mitochondrial function and injury responses, including potentially protective roles in ischemia/reperfusion and other stress-related settings [53].

Finally, overlapping loci near genes with vascular and endothelial relevance may indicate selection acting on pathways shaping vascular biology and fibrosis. The *MCAM* (*CD146*) gene is commonly described as an endothelial/vascular marker and has been linked to endothelial processes that intersect with fibrotic remodeling and vascular function, and it has also been explored as a biomarker or association signal in some cardiovascular studies [54,55]. Overall, the shared signals across ROH islands, iHS, and nSL suggest that selection in Doberman Pinschers may have targeted (or co-selected via linkage) genomic regions involved in ECM remodeling and cellular stress pathways, with additional contributions from vascular and neurodevelopment-related genes. These results provide coherent candidates for follow-up, including fine-mapping of overlapping intervals, deeper characterization of local haplotype structure, and integration with phenotype data (e.g., cardiac traits) to test whether these signals translate into measurable, breed-relevant outcomes.

### Interpopulation Selection Signatures Methods

To perform the interpopulation selection analyses, we compared the Doberman Pinscher population with a Labrador Retriever reference panel derived from Campbell et al. [25]. Labradors in this resource represent a well-characterized breed, making them a robust comparator across populations. This external reference allowed us to highlight Doberman-specific signals by directly comparing haplotype structure and allele-frequency patterns between breeds.

We applied three complementary approaches to capture population-specific selection in Dobermans. The FST (Fig 4A) quantifies allele-frequency differentiation between breeds; elevated FST values indicate loci that diverge more than expected under neutrality and may reflect breed-specific selection [56]. The XP-EHH (Fig 4B) contrasts the decay of haplotype homozygosity between populations and is particularly powerful for detecting completed or nearly completed sweeps, where selection has driven a haplotype close to fixation in one breed but not in the other [23]. The XP-nSL (Fig 4C) extends the nSL framework to a cross-population setting and leverages haplotype information while explicitly contrasting populations to detect both ongoing and near-fixed sweeps [24].

Because each metric emphasizes different aspects of the selective process, signals that agree across methods are more compelling than results from any single test. In general, FST reflects allele-frequency divergence, whereas XP-EHH and XP-nSL are more sensitive to sweep dynamics that are specific to one population, especially under strong selection when sweeps are advanced. Combining all three, therefore, provides a broader perspective on selection in the Doberman Pinscher genome across multiple evolutionary time scales. In the interpopulation analysis, we identified seven markers shared across methods (Fig 4D), mapping to regions containing 13 nearby genes. Several of these candidates are functionally consistent with pathways relevant to cellular regulation, neurobiology, and cardiovascular physiology. For example, the *BTG1* gene is involved in cell-cycle regulation and apoptosis and is widely studied in growth control [57], while the *ERMP1* gene is linked to endoplasmic-reticulum function and cellular stress responses and has also been discussed in neurobiological contexts [58,59]. The *NEGR1* gene has established roles in neuronal development and neurite arborization and has been associated with behavioral phenotypes, including anxiety- and aggression-related traits [60,61]. The *KCNK10* gene encodes a potassium channel, and variation in cardiac ion-channel genes has been implicated in canine dilated cardiomyopathy (DCM), including in Dobermans, which have high susceptibility to this condition [62–64]. In addition, the *SPATA7* gene encodes a ciliary scaffold protein required for the maintenance of the photoreceptor connecting cilium and proper protein trafficking in retinal cells [65], and the *PTPN21* gene encodes a non-receptor protein tyrosine phosphatase involved in intracellular signaling pathways that regulate cell growth and differentiation [66].

Overall, the cross-method overlap highlights a focused set of differentiated loci with consistent evidence for Doberman-specific selection and supports follow-up work to refine causal variants and connect these regions to breed-relevant phenotypes.

### Limitations and future directions

Certain methodological decisions in this study follow common practice for genome-wide selection scans, but there are also important limitations in how the findings should be interpreted and how candidate loci should be prioritized. Because selection statistics are continuous rather than discrete, researchers must define candidate regions using ranking-based criteria, which typically do not fit neatly into a formal hypothesis-testing framework. As a result, the specific cutoffs used to define candidate regions can meaningfully influence both the number and type of loci that are reported—especially for loci whose values fall close to the threshold.

In this study, candidate loci were identified using multiple outlier detection approaches. The ROH islands were defined as markers that occurred within an ROH in at least 50% of individuals. Haplotype-based candidates were defined using |iHS| and |nSL| > 2 for intrapopulation scans, and |XP-EHH| and |XP-nSL| ≥ 2 for cross-population scans. The ROH islands threshold used in this study has been previously reported in studies with dogs [67,68] and other populations as a practical criterion for defining ROH islands [69,70]. Regarding the other threshold, because the iHS, nSL, XP-EHH, and XP-nSL statistics are typically standardized across the genome — often within bins of allele frequency — score values are less dependent on underlying allele frequency. For that reason, a cutoff around |score| ≈ 2 is a commonly used heuristic cutoff for standardized haplotype signals [22–24]. However, the observed score distribution may not perfectly match test assumptions, so this type of cutoff should be interpreted as an outlier filter rather than a strict significance test [21,71,72].

For allele frequency differentiation, we used a stringent threshold (FST > 0.80) to prioritize markers that are highly differentiated between Doberman Pinschers and the Labrador population. This emphasizes extreme divergence consistent with near-fixation differences, but it may fail to capture more moderate patterns that could reflect polygenic selection and/or drift [73]. Finally, genotypes in this study were derived from SNP array data, which introduces uneven genome coverage and a bias towards common variants. Future work should validate key regions using imputed sequence data or whole-genome sequencing to help distinguish a single strong selective sweep from broader plateaus driven by linkage disequilibrium and/or overlapping selection signals.

## Conclusion

Overall, this study shows that the Doberman Pinscher population has high genome-wide autozygosity, consistent with recent inbreeding. Interestingly, the multiple approaches used for the selection signatures analysis showed convergent candidate signals within and between populations. These overlapping signals point to a broad functional process related to cellular modeling, stress, vascular and cardiac biology, with additional connections to neurodevelopmental processes and behavior. Overall, our findings provide a coherent set of pathways and genes that can be used as targets for follow-up studies integrating phenotypes relevant to breed health and performance.

## Acknowledgments

The authors would like to thank the Doberman Diversity Project for their valuable support and contribution to this study. We are especially grateful to Robin Loreth, Martina Fischer, and Carola Kusch, on behalf of the Doberman Diversity Project, for their assistance and collaboration throughout this work.

## Supporting information

**S1 Fig. Linkage disequilibrium decay (right) and principal components analysis (left) of the Doberman Pinscher population.**

**S1 Table. List of candidate genes near genomic markers commonly detected by three intrapopulation selection signature methods**

## References

1. Wade CM, Nuttall R, Liu S. Comprehensive analysis of geographic and breed-purpose influences on genetic diversity and inherited disease risk in the Doberman dog breed. Canine Med Genet. 2023;10: 7. doi:10.1186/s40575-023-00130-3

2. Meurs KM, Lahmers S, Keene BW, White SN, Oyama MA, Mauceli E, et al. A splice site mutation in a gene encoding for PDK4, a mitochondrial protein, is associated with the development of dilated cardiomyopathy in the Doberman pinscher. Hum Genet. 2012;131: 1319–1325. doi:10.1007/s00439-012-1158-2

3. Peťková B, Skurková L, Florian M, Slivková M, Dudra Kasičová Z, Kottferová J. Variations in Canine Behavioural Characteristics across Conventional Breed Clusters and Most Common Breed-Based Public Stereotypes. Animals. 2024;14. doi:10.3390/ani14182695

4. United Kennel Club. Breed Standards : Doberman Pinscher. 2019 [cited 4 Mar 2026]. Available: https://www.ukcdogs.com/doberman-pinscher?utm_source=chatgpt.com

5. Ivanovo UZUNOVA K, Stoycheva I, Miteva T, Bînev R, Ivanov A, Rousenov A, et al. Studies on Socialization Characteristics Using Two Temperament Tests in German Dogue, Doberman and Riesenschnautzer Puppies. Araştırma Makalesi J Fac Vet Med Istanbul Üniv. 2011;37: 43–51.

6. Meyers-Wallen V. Ethics and Genetic Selection in Purebred Dogs. Reproduction in Domestic Animals. 2003;38: 73–76. doi:10.1046/j.1439-0531.2003.00384.x

7. Peripolli E, Munari DP, Silva MVGB, Lima ALF, Irgang R, Baldi F. Runs of homozygosity: current knowledge and applications in livestock. Anim Genet. 2017;48: 255–271. doi:10.1111/age.12526

8. Mabunda RS, Makgahlela ML, Nephawe KA, Mtileni B. Evaluation of Genetic Diversity in Dog Breeds Using Pedigree and Molecular Analysis: A Review. Diversity (Basel). 2022;14. doi:10.3390/d14121054

9. Lewis TW, Abhayaratne BM, Blott SC. Trends in genetic diversity for all Kennel Club registered pedigree dog breeds. Canine Genet Epidemiol. 2015;2. doi:10.1186/s40575-015-0027-4

10. Hancock AM, Di Rienzo A. Detecting the genetic signature of natural selection in human populations: Models, methods, and data. Annu Rev Anthropol. 2008;37: 197–217. doi:10.1146/annurev.anthro.37.081407.085141

11. Oleksyk TK, Smith MW, O’Brien SJ. Genome-wide scans for footprints of natural selection. Philosophical Transactions of the Royal Society B: Biological Sciences. 2010;365: 185–205. doi:10.1098/rstb.2009.0219

12. Purcell S, Neale B, Todd-Brown K, Thomas L, Ferreira MAR, Bender D, et al. PLINK: A Tool Set for Whole-Genome Association and Population-Based Linkage Analyses. The American Journal of Human Genetics. 2007;81: 559–575. doi:10.1086/519795

13. VanRaden PM. Efficient Methods to Compute Genomic Predictions. J Dairy Sci. 2008;91: 4414–4423. doi:10.3168/jds.2007-0980

14. LI CC, HORVITZ DG. Some methods of estimating the inbreeding coefficient. Am J Hum Genet. 1953;5: 107. Available: https://pmc.ncbi.nlm.nih.gov/articles/PMC1716461/

15. Yang J, Lee SH, Goddard ME, Visscher PM. GCTA: A Tool for Genome-wide Complex Trait Analysis. The American Journal of Human Genetics. 2011;88: 76–82. doi:10.1016/j.ajhg.2010.11.011

16. Wright S. Coefficients of Inbreeding and Relationship. Am Nat. 1922;56: 330–338. doi:10.1086/279872

17. McQuillan R, Leutenegger A-L, Abdel-Rahman R, Franklin CS, Pericic M, Barac-Lauc L, et al. Runs of Homozygosity in European Populations. The American Journal of Human Genetics. 2008;83: 359–372. doi:10.1016/j.ajhg.2008.08.007

18. Pearson K. VII. Mathematical contributions to the theory of evolution.—III. Regression, heredity, and panmixia. Philosophical Transactions of the Royal Society of London Series A, Containing Papers of a Mathematical or Physical Character. 1896;187: 253–318. doi:10.1098/rsta.1896.0007

19. R Development Core Team. R: A language and environment for statistical computing. R Foundation for Statistical Computing. Vienna: Austria; 2009. Available: https://www.r-project.org/

20. Lencz T, Lambert C, DeRosse P, Burdick KE, Morgan TV, Kane JM, et al. Runs of homozygosity reveal highly penetrant recessive loci in schizophrenia. Proceedings of the National Academy of Sciences. 2007;104: 19942–19947. doi:10.1073/pnas.0710021104

21. Szpiech ZA, Hernandez RD. selscan: An Efficient Multithreaded Program to Perform EHH-Based Scans for Positive Selection. Mol Biol Evol. 2014;31: 2824–2827. doi:10.1093/molbev/msu211

22. Voight BF, Kudaravalli S, Wen X, Pritchard JK. A Map of Recent Positive Selection in the Human Genome. Hurst L, editor. PLoS Biol. 2006;4: e72. doi:10.1371/journal.pbio.0040072

23. Sabeti PC, Varilly P, Fry B, Lohmueller J, Hostetter E, Cotsapas C, et al. Genome-wide detection and characterization of positive selection in human populations. Nature 2007 449:7164. 2007;449: 913–918. doi:10.1038/nature06250

24. Szpiech ZA, Novak TE, Bailey NP, Stevison LS. Application of a novel haplotype-based scan for local adaptation to study high-altitude adaptation in rhesus macaques. Evol Lett. 2021;5: 408–421. doi:10.1002/evl3.232

25. Campbell CL, Bhérer C, Morrow BE, Boyko AR, Auton A. A Pedigree-Based Map of Recombination in the Domestic Dog Genome. G3 Genes|Genomes|Genetics. 2016;6: 3517–3524. doi:10.1534/g3.116.034678

26. Ayres DL, Darling A, Zwickl DJ, Beerli P, Holder MT, Lewis PO, et al. BEAGLE: An Application Programming Interface and High-Performance Computing Library for Statistical Phylogenetics. Syst Biol. 2012;61: 170–173. doi:10.1093/sysbio/syr100

27. Fonseca PAS, Suárez-Vega A, Marras G, Cánovas Á. GALLO: An R package for genomic annotation and integration of multiple data sources in livestock for positional candidate loci. Gigascience. 2020;9: 1–9. doi:10.1093/gigascience/giaa149

28. Dyer SC, Austine-Orimoloye O, Azov AG, Barba M, Barnes I, Barrera-Enriquez VP, et al. Ensembl 2025. Nucleic Acids Res. 2025;53: D948–D957. doi:10.1093/nar/gkae1071

29. Kolberg L, Raudvere U, Kuzmin I, Vilo J, Peterson H. gprofiler2 -- an R package for gene list functional enrichment analysis and namespace conversion toolset g:Profiler. F1000Res. 2020;9: 709. doi:10.12688/f1000research.24956.1

30. Wright S. THE INTERPRETATION OF POPULATION STRUCTURE BY F-STATISTICS WITH SPECIAL REGARD TO SYSTEMS OF MATING. Evolution (N Y). 1965;19: 395–420. doi:10.1111/j.1558-5646.1965.tb01731.x

31. Howrigan DP, Simonson MA, Keller MC. Detecting autozygosity through runs of homozygosity: A comparison of three autozygosity detection algorithms. BMC Genomics. 2011;12: 460. doi:10.1186/1471-2164-12-460

32. Mulim HA, Campos GS, Cardoso FF, Rojas de Oliveira H. Exploring inbreeding depression in Brazilian Angus cattle population using pedigree and genomic data. Front Genet. 2025;16. doi:10.3389/fgene.2025.1613820

33. Doekes HP, Bijma P, Windig JJ. How Depressing Is Inbreeding? A Meta-Analysis of 30 Years of Research on the Effects of Inbreeding in Livestock. Genes (Basel). 2021;12: 926. doi:10.3390/genes12060926

34. Villanueva B, Fernández A, Saura M, Caballero A, Fernández J, Morales-González E, et al. The value of genomic relationship matrices to estimate levels of inbreeding. Genetics Selection Evolution. 2021;53: 42. doi:10.1186/s12711-021-00635-0

35. Alemu SW, Kadri NK, Harland C, Faux P, Charlier C, Caballero A, et al. An evaluation of inbreeding measures using a whole-genome sequenced cattle pedigree. Heredity (Edinb). 2021;126: 410–423. doi:10.1038/s41437-020-00383-9

36. Tenhunen S, Thomasen JR, Sørensen LP, Berg P, Kargo M. Genomic analysis of inbreeding and coancestry in Nordic Jersey and Holstein dairy cattle populations. J Dairy Sci. 2024;107: 5897–5912. doi:10.3168/jds.2023-24553

37. Purfield DC, Berry DP, McParland S, Bradley DG. Runs of homozygosity and population history in cattle. BMC Genet. 2012;13: 70. doi:10.1186/1471-2156-13-70

38. Ferrer-Admetlla A, Liang M, Korneliussen T, Nielsen R. On Detecting Incomplete Soft or Hard Selective Sweeps Using Haplotype Structure. Mol Biol Evol. 2014;31: 1275–1291. doi:10.1093/molbev/msu077

39. Koumas L, King AE, Critchley HOD, Kelly RW, Phipps RP. Fibroblast Heterogeneity. Am J Pathol. 2001;159: 925–935. doi:10.1016/S0002-9440(10)61768-3

40. Pérez LA, Leyton L, Valdivia A. Thy-1 (CD90), Integrins and Syndecan 4 are Key Regulators of Skin Wound Healing. Front Cell Dev Biol. 2022;10: 810474. doi:10.3389/fcell.2022.810474

41. Rege TA, Hagood JS. Thy-1, a versatile modulator of signaling affecting cellular adhesion, proliferation, survival, and cytokine/growth factor responses. Biochimica et Biophysica Acta (BBA) - Molecular Cell Research. 2006;1763: 991–999. doi:10.1016/j.bbamcr.2006.08.008

42. Li Y, Song D, Mao L, Abraham DM, Bursac N. Lack of Thy1 defines a pathogenic fraction of cardiac fibroblasts in heart failure. Biomaterials. 2020;236: 119824. doi:10.1016/j.biomaterials.2020.119824

43. Glasson SS, Askew R, Sheppard B, Carito B, Blanchet T, Ma H-L, et al. Deletion of active ADAMTS5 prevents cartilage degradation in a murine model of osteoarthritis. Nature. 2005;434: 644–648. doi:10.1038/nature03369

44. Apte SS. Anti-ADAMTS5 monoclonal antibodies: implications for aggrecanase inhibition in osteoarthritis. Biochemical Journal. 2016;473: e1–e4. doi:10.1042/BJ20151072

45. Barallobre-Barreiro J, Radovits T, Fava M, Mayr U, Lin W-Y, Ermolaeva E, et al. Extracellular Matrix in Heart Failure: Role of ADAMTS5 in Proteoglycan Remodeling. Circulation. 2021;144: 2021–2034. doi:10.1161/CIRCULATIONAHA.121.055732

46. Delfín DA, DeAguero JL, McKown EN. The Extracellular Matrix Protein ABI3BP in Cardiovascular Health and Disease. Front Cardiovasc Med. 2019;6: 23. doi:10.3389/fcvm.2019.00023

47. Petersen JW, Douglas JY. Tenascin-X, collagen, and Ehlers–Danlos syndrome: Tenascin-X gene defects can protect against adverse cardiovascular events. Med Hypotheses. 2013;81: 443–447. doi:10.1016/j.mehy.2013.06.005

48. Matsumoto K, Aoki H. The Roles of Tenascins in Cardiovascular, Inflammatory, and Heritable Connective Tissue Diseases. Front Immunol. 2020;11: 609752. doi:10.3389/fimmu.2020.609752

49. Wagner MA, Siddiqui MAQ. The JAK-STAT pathway in hypertrophic stress signaling and genomic stress response. JAKSTAT. 2012;1: 131–141. doi:10.4161/jkst.20702

50. Wang S, Binder P, Fang Q, Wang Z, Xiao W, Liu W, et al. Endoplasmic reticulum stress in the heart: insights into mechanisms and drug targets. Br J Pharmacol. 2018;175: 1293–1304. doi:10.1111/bph.13888

51. Xing J, Li P, Hong J, Wang M, Liu Y, Gao Y, et al. Overexpression of Ubiquitin-Specific Protease 2 (USP2) in the Heart Suppressed Pressure Overload-Induced Cardiac Remodeling. Mediators Inflamm. 2020;2020: 1–12. doi:10.1155/2020/4121750

52. Rafiq K, Kolpakov MA, Seqqat R, Guo J, Guo X, Qi Z, et al. c-Cbl Inhibition Improves Cardiac Function and Survival in Response to Myocardial Ischemia. Circulation. 2014;129: 2031–2043. doi:10.1161/CIRCULATIONAHA.113.007004

53. Xiao Y, Rudolphi C, Nollet E, Bakker D, Nederlof R, Wang Q, et al. The innate immune receptor NLRX1 protects against cardiac ischemia/reperfusion injury through RISK signaling, mPTP regulation and inhibition of mitochondrial activity. J Mol Cell Cardiol. 2022;173: S8–S10. doi:10.1016/j.yjmcc.2022.08.020

54. Mocan D, Jipa R, Jipa DA, Lala RI, Puschita M, Rasinar F-C, et al. Soluble CD146 in Heart Failure: Pathophysiological Role and Diagnostic Potential. Biomedicines. 2025;13: 1370. doi:10.3390/biomedicines13061370

55. Zhang Z-Y, Zhai C, Yang X-Y, Li H-B, Wu L-L, Li L. Knockdown of CD146 promotes endothelial-to-mesenchymal transition via Wnt/β-catenin pathway. Chadjichristos CE, editor. PLoS One. 2022;17: e0273542. doi:10.1371/journal.pone.0273542

56. Boca SM, Rosenberg NA. Mathematical properties of Fst between admixed populations and their parental source populations. Theor Popul Biol. 2011;80: 208–216. doi:10.1016/j.tpb.2011.05.003

57. Yuniati L, Scheijen B, van der Meer LT, van Leeuwen FN. Tumor suppressors BTG1 and BTG2: Beyond growth control. J Cell Physiol. 2019;234: 5379–5389. doi:10.1002/jcp.27407

58. Solarz-Andrzejewska A, Majcher-Maślanka I, Kryst J, Chocyk A. Modulation of the endoplasmic reticulum stress and unfolded protein response mitigates the behavioral effects of early-life stress. Pharmacological Reports. 2023;75: 293–319. doi:10.1007/s43440-023-00456-6

59. Grandi A, Santi A, Campagnoli S, Parri M, De Camilli E, Song C, et al. ERMP1, a novel potential oncogene involved in UPR and oxidative stress defense, is highly expressed in human cancer. Oncotarget. 2016;7: 63596–63610. doi:10.18632/oncotarget.11550

60. Pischedda F, Piccoli G. The IgLON Family Member Negr1 Promotes Neuronal Arborization Acting as Soluble Factor via FGFR2. Front Mol Neurosci. 2016;8: 175299. doi:10.3389/fnmol.2015.00089

61. Singh K, Loreth D, Pöttker B, Hefti K, Innos J, Schwald K, et al. Neuronal Growth and Behavioral Alterations in Mice Deficient for the Psychiatric Disease-Associated Negr1 Gene. Front Mol Neurosci. 2018;11. doi:10.3389/fnmol.2018.00030

62. Polenberg A, Grueter C, Braun T, Mitchell C. A region on chromosome 16 is associated with Doberman Pinscher dilated cardiomyopathy. Sci Rep. 2024;14: 27241. doi:10.1038/s41598-024-78511-2

63. Simpson S, Edwards J, Ferguson-Mignan TFN, Cobb M, Mongan NP, Rutland CS. Genetics of Human and Canine Dilated Cardiomyopathy. Int J Genomics. 2015;2015: 1–13. doi:10.1155/2015/204823

64. Talley EM, Solórzano G, Lei Q, Kim D, Bayliss DA. CNS Distribution of Members of the Two-Pore-Domain (KCNK) Potassium Channel Family. The Journal of Neuroscience. 2001;21: 7491–7505. doi:10.1523/JNEUROSCI.21-19-07491.2001

65. Eblimit A, Nguyen T-MT, Chen Y, Esteve-Rudd J, Zhong H, Letteboer S, et al. Spata7 is a retinal ciliopathy gene critical for correct RPGRIP1 localization and protein trafficking in the retina. Hum Mol Genet. 2015;24: 1584–1601. doi:10.1093/hmg/ddu573

66. Cho Y-C, Kim BR, Cho S. Protein tyrosine phosphatase PTPN21 acts as a negative regulator of ICAM-1 by dephosphorylating IKKβ in TNF-α-stimulated human keratinocytes. BMB Rep. 2017;50: 584–589. doi:10.5483/BMBRep.2017.50.11.169

67. Huson HJ, Srikanth K, Ellis KM. Breeding Selection for U.S. Siberian Huskies Has Altered Genes Regulating Metabolism, Endurance, Development, Body Conformation, Immune Function, and Behavior. Genes (Basel). 2025;16: 1355. doi:10.3390/genes16111355

68. Sweetalana, Nataneli S, Huang S, Mooney JA, Szpiech ZA. Genotypic and phenotypic consequences of domestication in dogs. bioRxiv. 2025; 2024.05.01.592072. doi:10.1101/2024.05.01.592072

69. Mulim HA, Brito LF, Batista Pinto LF, Moletta JL, Da Silva LR, Pedrosa VB. Genetic and Genomic Characterization of a New Beef Cattle Composite Breed (Purunã) Developed for Production in Pasture-Based Systems. Front Genet. 2022;13. doi:10.3389/fgene.2022.858970

70. Peripolli E, Stafuzza NB, Amorim ST, de Lemos MVA, Grigoletto L, Kluska S, et al. Genome-wide scan for runs of homozygosity in the composite Montana Tropical® beef cattle. Journal of Animal Breeding and Genetics. 2020;137: 155–165. doi:10.1111/jbg.12428

71. Yao Y, Pan Z, Di R, Liu Q, Hu W, Guo X, et al. Whole Genome Sequencing Reveals the Effects of Recent Artificial Selection on Litter Size of Bamei Mutton Sheep. Animals 2021, Vol 11, Page 157. 2021;11: 157. doi:10.3390/ani11010157

72. Li X, Yang J, Shen M, Xie XL, Liu GJ, Xu YX, et al. Whole-genome resequencing of wild and domestic sheep identifies genes associated with morphological and agronomic traits. Nat Commun. 2020;11: 2815. doi:10.1038/s41467-020-16485-1

73. Buckley RM, Bilgen N, Harris AC, Savolainen P, Tepeli C, Erdoğan M, et al. Analysis of canine gene constraint identifies new variants for orofacial clefts and stature. Genome Res. 2025;35: 1080–1093. doi:10.1101/gr.280092.124

